# The proteome content of blood clots observed under different conditions: successful role in predicting clot amyloid(ogenicity)

**DOI:** 10.1101/2024.11.29.626062

**Authors:** Douglas B. Kell, Etheresia Pretorius

## Abstract

A recent analysis compared the proteome of (i) blood clots seen in two diseases – sepsis and long COVID – when blood was known to have clotted into an amyloid microclot form (as judged by staining with the fluorogenic amyloid stain thioflavin T) with (ii) that of those non-amyloid clots considered to have formed normally. Such fibrinaloid microclots are also relatively resistant to fibrinolysis. The proteins that the amyloid microclots contained differed markedly both from the soluble proteome of typical plasma and that of normal clots, and also between the disease studies (an acute syndrome in the form of sepsis in an ITU and a chronic disease represented by Long COVID). Many proteins in the amyloid microclots were low in concentration in plasma and were effectively accumulated into the fibres, whereas many other abundant plasma proteins were excluded. The proteins found in the microclots associated with the diseases also tended to be themselves amyloidogenic. We here ask effectively the inverse question. This is: can the clot proteome tell us whether the clots associated with a particular disease contained proteins that are observed uniquely (or are highly over-represented) in known amyloid clots relative to normal clots, and thus were in fact amyloid in nature? The answer is in the affirmative in a variety of major coagulopathies, viz. venous thromboembolism, pulmonary embolism, deep vein thrombosis, various cardiac issues, and ischaemic stroke. Galectin-3-binding protein and thrombospondin-1 seem to be especially widely associated with amyloid-type clots, and the latter has indeed been shown to be incorporated into growing fibrin fibres. These may consequently provide useful biomarkers with a mechanistic basis.

## 1. Introduction

Blood normally clots into a thrombus in which the fibres appear like cooked spaghetti when observed in the electron microscope (e.g. [1-5]). However, a series of our own studies [6-23], as well as those of others [24-26], have demonstrated, via staining with the amyloid stain thioflavin T (or by other, label-free means [27]), that fibrinogen monomers (when exposed to thrombin) can self-assemble to form an amyloid version of (micro)clots that we refer to as fibrinaloid microclots. As with other forms of amyloid proteins, these are much more resistant to proteolysis (fibrinolysis [28]) than are the normal forms. In a recent analysis [29], we compared the proteome of blood clots seen when blood had clotted into a known amyloid form (as judged by staining with the classical amyloid stain thioflavin T) with those clots considered to have formed normally. While the amounts of non-fibrin proteins that were trapped in the normal clots had a rough correlation with the typical contents of the plasma proteome, there was little such correlation in the amyloid clots; indeed the proteins that the amyloid microclots contained were characterised by their highly amyloidogenic potential [29]. They also differed markedly between the disease studies (viz acute sepsis in an ITU [25] vs the chronic disease Long COVID [9], with only apolipoprotein-A2 being common to both). Notably, we there showed that the fibrinaloid microclots are associated with proteins that are very different [9,25] from those within normal, non-amyloid clots, using for the latter data from a paper by Undas and colleagues [30]. More explicitly, we argued there [29] that, in the terminology of Bondarev and colleagues [31], as illustrated in Figure 1, proteins entrapped in normal clots were likely to be ‘titrated’ (bound) or sequestered, while those in fibrinaloid microclots were, through their cross-β motifs, likely to be incorporated such as to be part of the amyloid fibres themselves. All of this, along with the presence of anti-fibrinolytic proteins such as α2-antiplasmin [28], served to explain the significant resistance of these microclots to normal fibrinolysis. As it happens, there are a number of other clot proteomics studies available, studying samples from a variety of diseases (e.g. [32,33]), but in which the amyloidogenic nature of the clots was not, however, measured. Since we now have access to proteomes from known, ‘calibrated’ systems, we can therefore here effectively ask the inverse question (Figure 2). This is: can the clot proteome tell us whether the clots associated with a particular disease plausibly contained proteins that are observed uniquely (or are highly over-represented) in known amyloid clots relative to normal clots, and thus that the clots were in fact probably amyloid in nature? The present paper sets out to answer this; in all cases where the data are available, the answer is in the affirmative, suggesting the value of testing this directly, as has very recently been done in the case of clots removed by thrombectomy from stroke patients [34].

**Figure 1:**
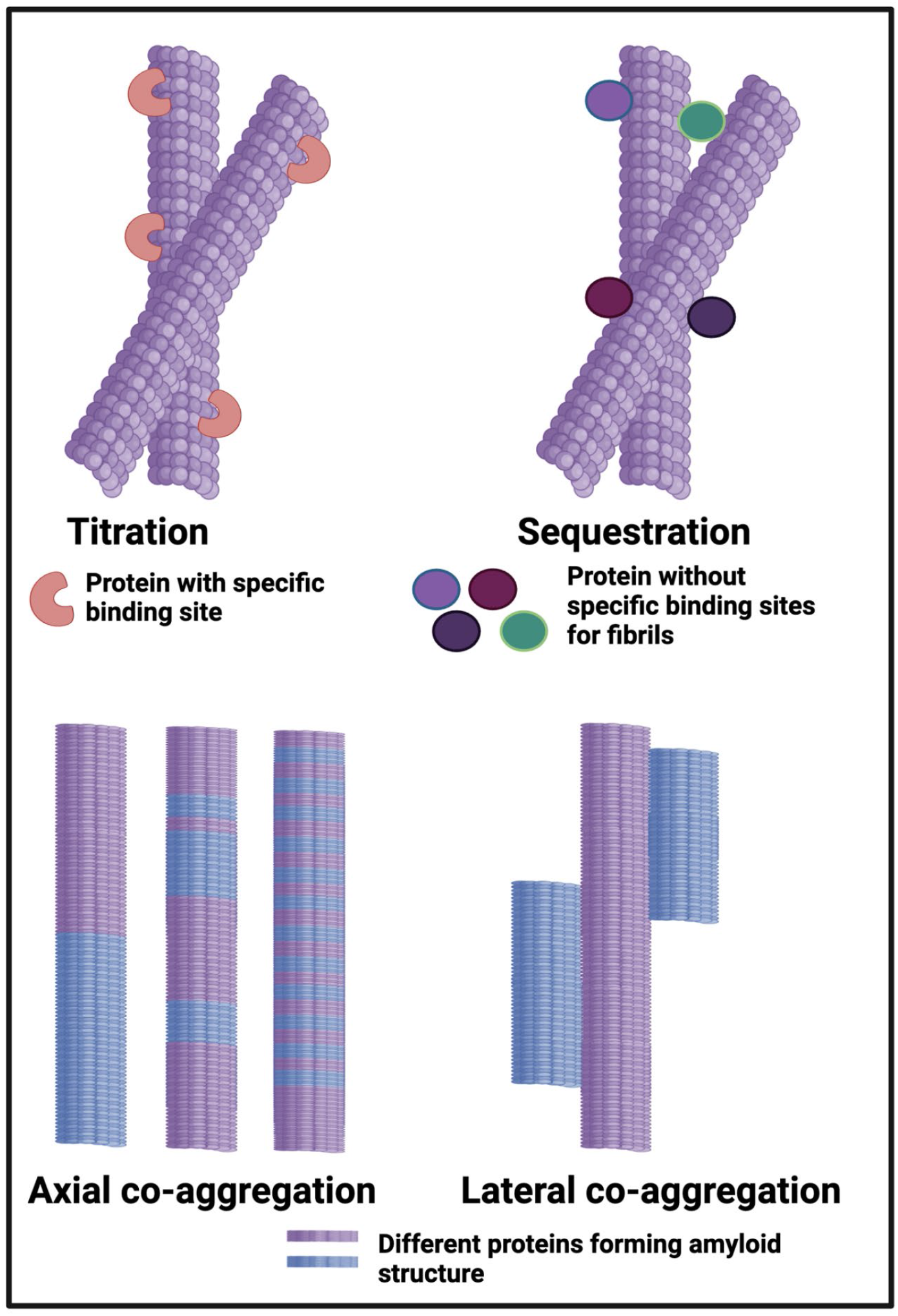
Different classes or types of protein co-aggregation: **(A)** Titration; (**B)** Sequestration; **(C)** Axial and **(D)** Lateral. Reprinted from the open access preprint [29], which was itself adapted from [31].

**Figure 2.**
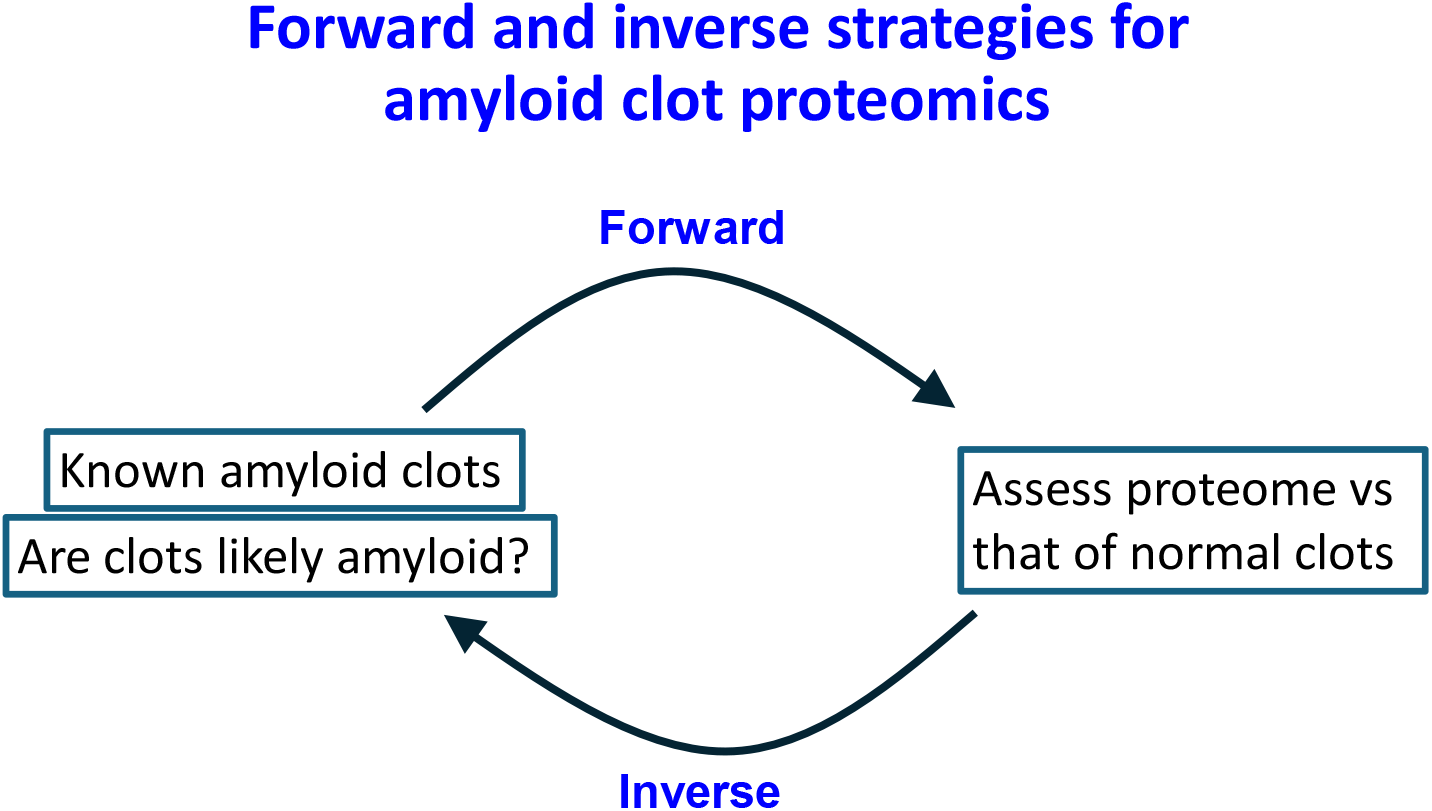
The relationship between forward and inverse strategies (as in systems biology [35]), in which here we seek to assess the normal or amyloid nature of clots as judged by their proteome. In the forward strategy we calibrate the system by asking which proteins differ in the two cases where the amyloid nature is known [36]. In the inverse case, developed here, we use the observed protein entrapments to infer or to suggest whether the clots are likely to be amyloid in nature.

## 2. Results

Letunica and colleagues [32] provide a very helpful summary of clot proteome studies, and in what follows we analyse the relevant (clot) studies that they highlight, along with some other studues of clot proteomes under various circumstances.

Alonso-Orgaz and colleagues [37] studied the proteome of clots taken from individuals who had suffered an acute myocardial infarction with ST-segment elevation (STEMI patients). They used three mass spectrometric methods for proteome analysis; we here confine ourselves to the 46 proteins detected using all three methods (their table S5). In fact, apart from fibrin(ogen), there was surprisingly little overlap with any of the proteins in the previous works [9,25,30] that we studied, with the exception of thrombospondin-1 [37]; however, thrombospondin-1 (Uniprot P07996) was present in our Long COVID study [9] but barely so (8 μg.g^-1^ clot) in the normal-clot study of Undas and colleagues [30]). This molecules thus provides a strong hint for the amyloid nature of the clots in the STEMI patients.

### Venous thromboembolism (VTE)

Stachowicz and colleagues [38] studied the proteome of clots generated from the plasma of individuals with venous thromboembolism (VTE). Interestingly, these clots were significantly resistant to trypsinolysis (they yielded about 4x more peptides using lys C), possibly providing a strong hint (as per [39]) that these clots were likely amyloid in character, albeit this was not measured. LysC + trypsin + chymotrypsin yielded the most peptides [38] (note that in *E. coli* the peptides yielded by incubation with lys C vs trypsin were largely orthogonal [40]). Of the proteins mentioned in their Table 1 [38], fibronectin, vitronectin and antithrombin III are more typical of proteins entrapped in normal clots, while α2-antiplasmin and von Willebrand factor appear in both [29].

VTE was also studied by Bruzelius and colleagues [41]. The stand-out markers for VTE in their study were VWF and platelet-derived growth factor B (PDGFB); the latter did not make an appearance in any of our earlier studies. VWF is also very prone to changes in that it may be elevated at one stage in a coagulopathic disease’s progression but hypercoagulability can deplete it rapidly, even thereby leading to hypocoagulability [42].

Another prospective VTE study was that of Jensen and colleagues [43], who analysed the proteome of those suffering a VTE in samples obtained from a large cohort study. Most interestingly, one of their top biomarkers was Galectin-3-binding protein (LG3P), a protein that was highly over-represented in our Long COVID study of fibrinaloid microclots [9], but that did not appear in the normal clot proteome [30]. Transthyretin, a protein well known to be prone to amyloidogenesis (e.g. [44-46], was also upregulated in those clot proteomes whose owners later went on to have a VTE [43]. Galectin-3-binding protein (LG3P) does seem like an interesting marker for <u>later</u> VTE events, and is certainly amyloidogenic.

### Pulmonary embolism

Bryk and colleagues [47] studied the quantitative proteome of clots generated from the plasma of 20 individuals who had had a pulmonary embolism (PE) compared with that of 20 controls. They [47] used a combination of lysC and trypsin to ensure good digestion. The specific protein composition in plasma fibrin clots from acute PE patients was associated with denser clots, and proteins that were enriched in these included apolipoprotein B, a protein enriched in our own studies [9] of fibrinaloid microclots. PE is hugely exacerbated by COVID-19 [48], so it is probably not then surprising (e.g. [9,17,18,20,39,42,49-59]) to see amyloid-type clot markers in these clots. Bryk and colleagues also detected platelet factor 4 in their clots, a protein also detected in fibrinaloid microclots by Schofield *et al*. [25], but present only in minuscule amounts in both plasma and normal clots [29]. These findings suggest strongly that the clots seen with pulmonary embolism are also indeed likely to be amyloid in nature.

In an important series of studies Ząbczyk *et al*. [5,60-62] analysed the ‘prothrombotic fibrin clot phenotype’ that they discovered to be associated with pulmonary embolisms. Recurrent PE was associated with the formation of denser fibrin networks reflected by lower permeability, with impaired fibrinolysis, and consequently with reduced maximum rate of increase in D-dimer levels in the lysis assay, despite higher plasma D-dimer levels [62]. Although seemingly not measured, these are precisely the properties of fibrinaloid microclots. The prothrombotic fibrin clot properties were associated with NETs formation [5,61] (see also [63]) as well as elevated lactate levels [61] (implying hypoxia), and most importantly of all (from our perspective) with low grade endotoxaemia [60]. Endotoxin (bacterial lipopolysaccharide) is <u>precisely</u> the trigger that we initially discovered [11], and have subsequently confirmed many times [13-16,64], that was able to catalyse the formation of fibrinaloid microclots. It is hard not to infer that these clots are amyloid in nature; hopefully those studying them will deploy the necessary tests, which are easy enough to do (e.g. [65,66]).

### Deep vein thrombosis (DVT)

As a subset of VTE, Deep Vein Thrombosis (DVT) is seen as less likely to be fatal than is PE, but is nonetheless a significant cardiovascular problem. In addition, post-thrombotic syndrome (PTS) is a common comorbidity (of DVT). We suspect that the two are related via fibrinaloid microclots, since the main clinical manifestation of PTS is chronic venous insufficiency [67,68]. Ramacciotti and colleagues studied the proteome of microparticles following DVT. Very strikingly just two proteins, namely Galectin-3 Binding Protein (LG3BP) and Alpha-2 macroglobulin (a highly expressed plasma protein [69]) were over-expressed. As noted [29], the amyloidogenic LG3BP is highly over-represented in the fibrinaloid microclots of Long COVID, but is invisible in normal clots. Zhang and colleagues studied the plasma (rather than clot) proteome of individuals with various kinds of DVT [70], and again found a great many differences (Figure 3), in addition suggesting several metabolites that might be discriminatory. Interestingly the protein most lowered <u>in the plasma</u> of patients showing DVT following trauma (pt-DVT) (Figure 3) was the highly amyloidogenic [44,71] transthyretin, strongly implying that it had disappeared into amyloidogenic clots.

**Figure 3.**
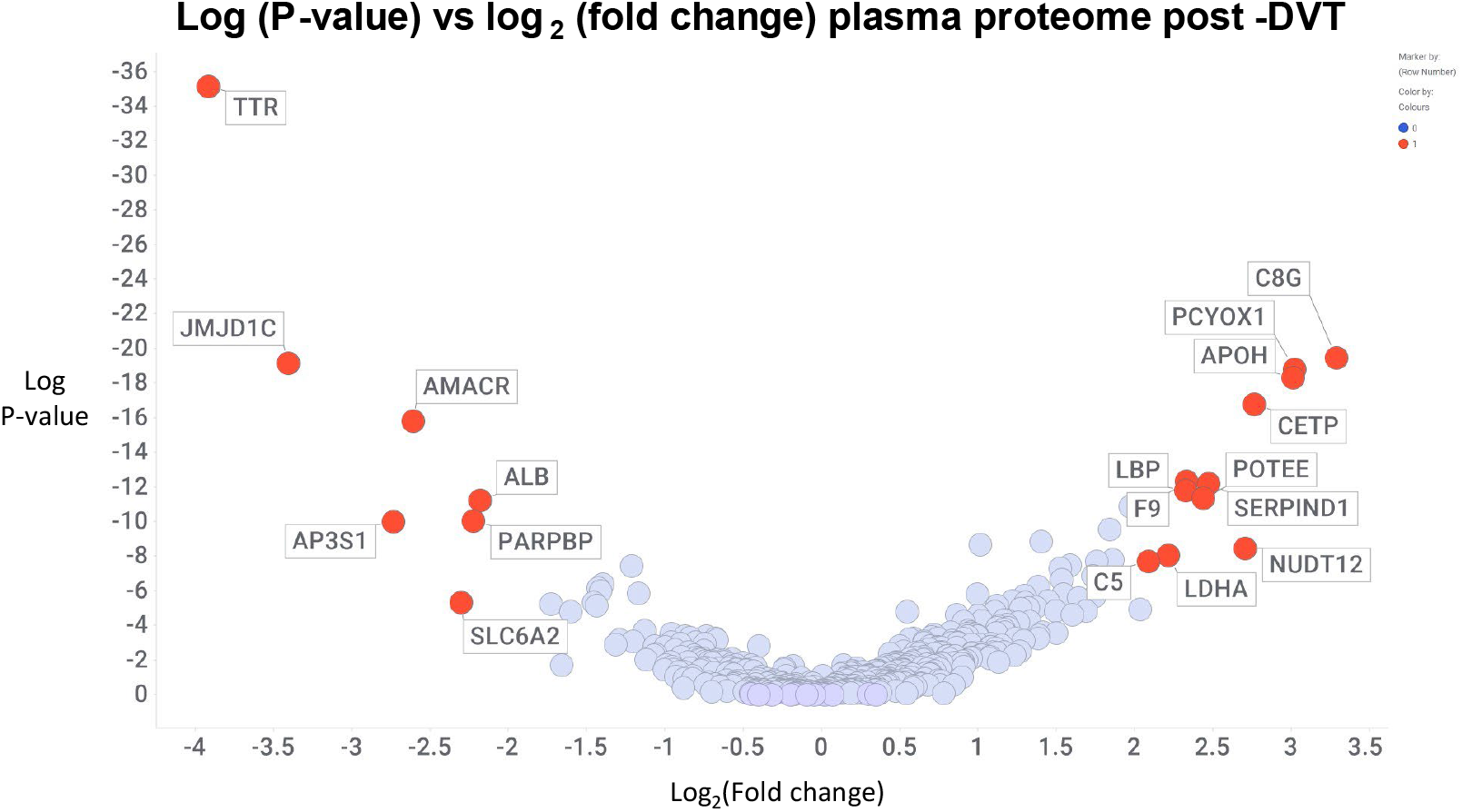
Changes in the plasma proteome in individuals with deep vein thrombosis after trauma. Data are taken from supplementary table 15 of [70] and visualised using the Spotfire program (http://spot-fire.com/). Those with a log2 change <-2 or >2 are labelled

### Post-thrombotic syndrome

Post thrombotic syndrome (PTS) is a common chronic complication of deep vein thrombosis [72], occurring in up to 50% of individuals who have had a DVT [73]. It seems to involve residual clots that are not removed, in that successful therapies include catheter-directed thrombolysis and mechanical thrombectomy [73,74], as well as low-MW heparin [75]. Obviously we suspect fibrinaloid clots here, though no one seems to have looked, and part of the purpose of this article is to encourage experts to do so.

### Cardioembolic and large artery atherosclerotic clots

Rossi and colleagues [76] studied the proteome of formalin-fixed paraffin-embedded cardioembolic and large artery atherosclerotic (LAA) clots, finding 1,581 proteins; however, the numbers of samples were insufficient for us to draw meaningful conclusions.

### Myocardial infarction

We have been unable to find any studies in which clots were analysed following a myocardial infarction (MI). However, it is notable that serum amyloid A is a risk factor for MI [77] and is massively increased in patients who have had an acute myocardial infarction [78]. It is to be stressed that serum amyloid A has been shown to promote fibrinaloid formation [10].

### Trauma-induced coagulopathy

A somewhat separate but closely related category is trauma-induced coagulopathy (TIC). This describes abnormal coagulation processes that are attributable to trauma [79]. In contrast to acute COVID, in which the sequence is opposite [42], TIC exhibits a hypocoagulable phase linked to haemorrhage [79-81] followed by one of hypercoagulability [79,82]. Both seem to be controlled significantly by the rates of fibrinolysis [83,84], with the hypocoagulability being attributed to hyperfibrinolysis, and the hypercoagulability to hypofibrinolysis. For present purposes, our focus is on the latter, with hypofibrinolysis being explicable, at least in part, by any amyloid formation. As per the theme of this article, we next explore what is known of the proteome in the hypercoagulable phase of TIC.

Sadly, such studies of the clot proteome in TIC seem few and far between. However, Coleman and colleagues [85] found increases in both α2-antiplasmin and α1-antitrypsin (SERPINA1), both of which are increased in fibrinaloid microclots [9,36,39], are stimulated by oestradiol (that may also contribute to this phenotype [86]) and that this can contribute to the known sex dimorphism in coagulability [87]. Interestingly, thrombospondin-1 was among the most overexpressed proteins (3.84-fold; supplementary information to [85]) and this is one of our two most favoured markers for clots being amyloid in nature [36]. Apolipoprotein-A1 was also strongly over-represented in the clot proteomes of [85], and this too was a significant contributor to the amyloid clot proteome of Toh and colleagues [25]. Taken together, the evidence strongly favours the view that the clots produced when a patient is exhibiting hypercoagulability in TIC are indeed amyloid in nature.

### Stroke

Stroke, the second greatest cause of death worldwide [88-91], has long been associated with the production of anomalous clots [92-94] (that, as rehearsed above, we now know to be amyloid in nature).

O’Connell and colleagues [95] studied the circulating (rather than clot) proteome of individuals following an ischaemic stroke. They found a substantial gender difference in corticosteroid-binding protein that rather dominated the analyses and overwhelmed stroke effects.

Penn et al sought proteomic biomarkers of stroke. Their first study of the clot proteome [96] found proteins involved in inflammation (47%), coagulation (40%), and atrial fibrillation (7%), among others. Atrial fibrillation is, of course, well associated with stroke risk [97], as well as a prothrombotic state [98-100]. Other studies known as SpecTRA [101,102] included the assessment of thrombospondin and VWF as part of a biomarker panel. Further validation studies showed that both thrombospondin and apolipoprotein B contributed. Both have been recognised as amyloidogenic markers in fibrinaloid microclots, and in particular it is to be highlighted that thrombospondin can be incorporated into fibrin fibrils themselves [103-106] (and see later).

Lopez-Pedrera and colleagues [107] analysed the thrombus proteome of stroke patients. The thrombus proteome reflected three classes or clusters of patients with varying levels of severity, prognosis, and aetiology of the stroke. Gelsolin and vinculin were among the proteins observed in the clots. Cardioembolic thrombi were enriched in proteins of the innate immune system.

Staessens and colleagues [108,109] note the prevalence of neutrophil extracellular traps (NETs) in the thrombus taken from ischaemic strokes (they were also found in cardioembolic thrombi [110]); very interestingly NETs are also an important feature of the fibrini-aloid microclots found in long COVID [51], and were found in prothrombotic clots in the studies of Ząbczyk *et al*. [5]. Lower levels of extracellular DNA were related to the ease of thrombectomy [111], plausibly allowing one to relate the extent of amyloid clotting to the difficulty of thrombectomy (often measured as the number of passes).

Muñoz and colleagues [112] analysed the proteome content of clots from ischaemic strokes, noting some 1600 proteins, but quantitative data were not given and so we cannot extract meaningful conclusions for the present purposes.

Finally, Prochazka and colleagues [113] analysed clots following acute ischaemic stroke, noting the role of von Willebrand Factor and ADAMTS13.

Most pertinently, however, is that we have shown experimentally, and for the first time, that the clots thrombectomised from 8/8 individuals with ischaemic stroke were strongly amyloid in character [34], strongly validating the arguments put forward here.

Two of the proteins over-represented in fibrinaloid microclots that regularly appeared in coagulopathic diseases are illustrated in Figure 4.

**Figure 4:**
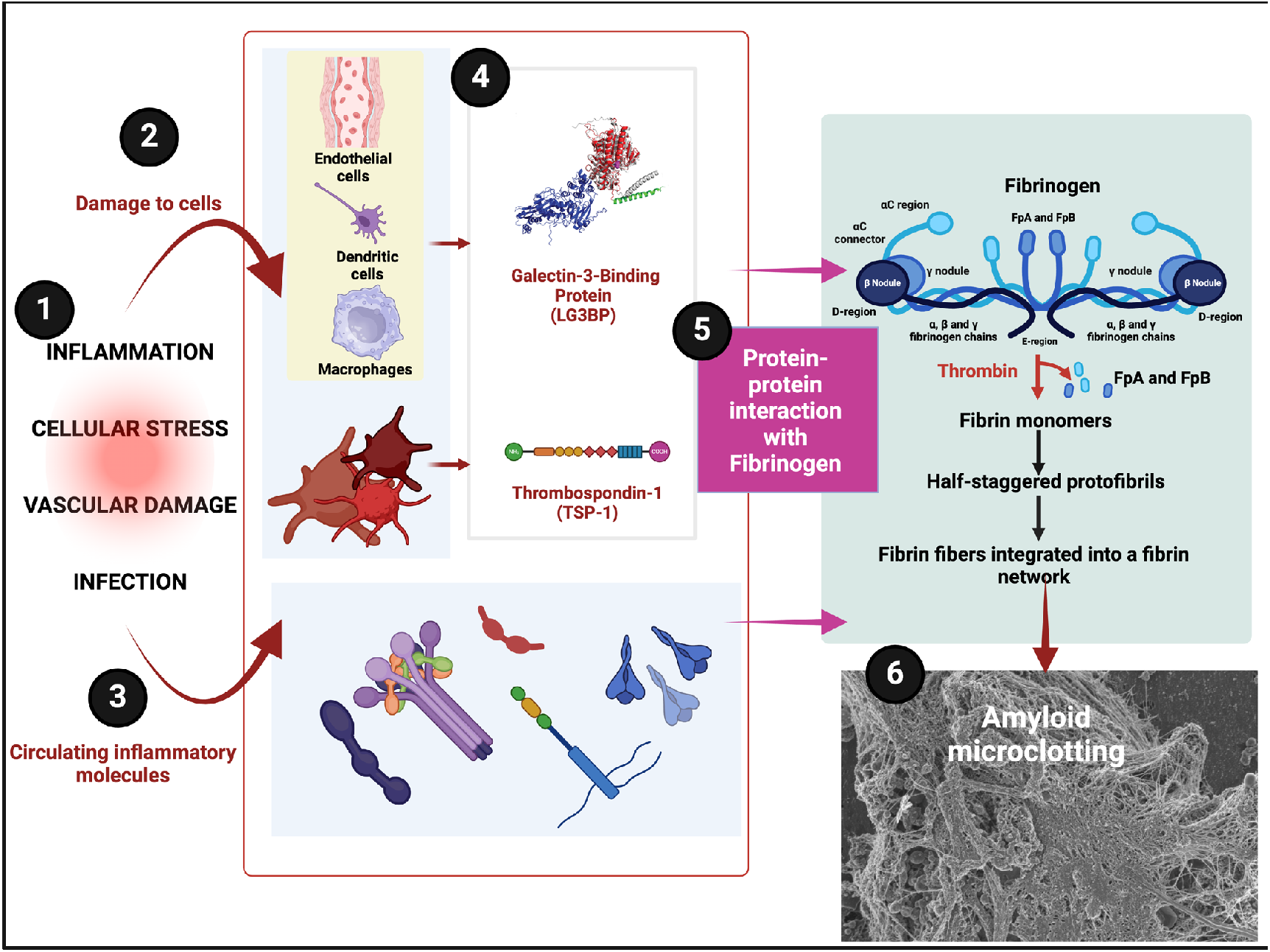
Illustration of the pathways from inflammation-induced cellular damage to the formation of amyloid microclots, highlighting the roles of two notably amyloidogenic proteins (LG3BP and TSP-1) in abnormal fibrinogen interactions and the resulting fibrinolysis-resistant clot structures. **1)** Inflammation and Cellular Stress: Triggers that initiate vascular damage, often due to infection or systemic inflammation. **2)** Damage to Cells: Injury to endothelial cells and activation of immune cells like dendritic cells and macrophages. **3)** Circulating Inflammatory Molecules: Release of inflammatory mediators that exacerbate vascular stress and promote clot formation. **4)** Key Amyloidogenic Proteins: Production of Galectin-3-Binding Protein (LG3BP) and Thrombospondin-1 (TSP-1) by immune and endothelial cells. **5)** Protein-Protein Interaction with Fibrinogen: Interaction of LG3BP and TSP-1 with fibrinogen, promoting amyloid-like fibrin formation. **6)** Amyloid Microclotting: Formation of amyloid microclots that are resistant to fibrinolysis, contributing to disease pathology.

### Galectin-3-Binding Protein (LG3BP)

LG3BP (Uniprot B4DVE1), also known as Mac-2 binding protein (Mac-2BP) or tumor-associated antigen 90K [114], is a heavily glycosylated, secreted protein whose expression is induced in viral infection and other inflammatory conditions [115-117], including metabolic syndrome [118,119] and cancer [120,121]. LG3BP appears in so many examples of the clots seen in a variety of the coagulopathic diseases under study, as well as in established fibrinaloid microclots in Long COVID [9], that it seems beyond peradventure for it not to have an aetiological role. It does not seem to be present in normal clots [30]. It has already been highlighted as a major player in the development of deep vein thrombosis [122-125], albeit detailed mechanisms are not to hand. However, the fact that it can participate in fibrinaloid microclots plausibly means that it can induce them. Certainly it is part of the deposits seen in glomerular nephritis [126], but these seemingly have not been tested for amyloid properties either. This said, renal amyloidosis is a common accompaniment to glomerulonephritis [127] and kidney damage more generally [128-133]. Galectin-3, the binding partner of LG3BP, is itself amyloidogenic [134], an inducer of fibrosis [135-138], and is a risk factor for for a variety of diseases [138-140] including atrial fibrillation [141-145], something highly pertinent to this discussion [97].

Given the propensity for cross-seeding (many references, summarised in [29]), it is noteworthy that L3GBP is also capable of interacting with the precursor of the highly amyloidogenic Aβ protein [146].

### Thrombospondin-1

Thrombospondin-1 (TSP-1) (Uniprot P07996) is a homotrimeric, heavily glycosylated [147], 450 kDa glycoprotein stored within the α-granules of platelets and released upon platelet activation [148]. It has significant roles in a variety of cardiovascular diseases [149-152] (including atrial arrhythmias [153]; see also [97]), lung pathologies [154], carcinomas [155,156], ageing [157], and, interestingly, stimulating platelet aggregation [158] and fibrosis [159-163], though it remains comparatively under-studied [164]. It is known to interact with Aβ [165], and, most importantly for the present analysis, it was shown nearly 40 years ago [103,105,106] that it is actually incorporated into fibrin during clot formation. Although this was not measured at the time, it seems likely that those clots were amyloid in nature, especially since – as with known fibrinaloid microclots [11] – the fibres so formed were on average both thinner and more numerous than clots formed in its absence [104].

Although thrombospondin-1 levels are comparatively low in platelet-poor plasma [148], thrombospondin-1 was highly enriched in the fibrinaloid microclots associated with Long COVID [9] but barely detectable in normal clots [30]. It seems that in clot proteomics thrombospondin-1, along with galectin-3-binding protein, could indeed be an excellent marker of fibrinaloids, and may even induce them.

## 3. Discussion and conclusions

In a previous analysis [29], we assessed the differences in the clot proteome between clots known to be normal and those measured experimentally to be amyloid in character (fibrinaloid clots), concluding that the major difference in those proteins that were over- or under-represented in the clots was clearly correlated with the extent to which they were amyloidogenic in character. Coupled with the contrasting fact that the proteome content of normal clots was far more representative of the normal plasma proteome, this gave strong weight to the view that the ‘entrapment’ of the non-fibrin proteins was actually within the cross-β elements of the amyloid fibrils themselves, via so-called cross-seeding.

In the present analysis we asked what amounts to the inverse question, which is as follows: given what we know about proteins enriched in amyloid clots, could we seek to <u>predict</u> the amyloid nature of the clots in a variety of diseases in which it was not measured by looking at the proteome alone? The answer was a resounding yes. In particular, galectin-3-binding-protein and thrombospondin-1 seemed to be present in all the clots from the diseases studied here, and both were highly amyloidogenic (scoring over 0.86 on the amylogram server [166,167] http://biongram.biotech.uni.wroc.pl/AmyloGram/). It now seems that the polymerisation of fibrin(ogen) into an amyloid is indeed associated with a considerable number of acute, as well as chronic, diseases.

## 4. Materials and methods

This was an analytical paper; the data were obtained and the analyses performed precisely as described in the text.

## Supplementary Materials

Not applicable.

## Author Contributions

Both authors contributed to the conceptualisation, analyses, funding acquisition, drafting, and final editing.

## Acknowledgments

DBK thanks Prof Ben Goult for useful discussions.

## Disclaimer/Publisher’s Note

The statements, opinions and data contained in all publications are solely those of the individual author(s) and contributor(s) and not of MDPI and/or the editor(s). MDPI and/or the editor(s) disclaim responsibility for any injury to people or property resulting from any ideas, methods, instructions or products referred to in the content.

